# Strategies to improve the efficiency of homing gene drives with multiplexed gRNAs

**DOI:** 10.1101/2025.06.28.662113

**Authors:** Weizhe Chen, Peixin Wu, Jackson Champer

## Abstract

CRISPR homing gene drive holds great potential for pest control, but its success is challenged by the generation of resistance alleles. To mitigate the impact of resistance, multiplexed gRNA strategies have been demonstrated. However, unless both outmost sites are cleaved simultaneously, poor homology during DNA repair may compromise efficiency, leading to decline in drive conversion efficiency when the number of gRNAs is higher than two. Here, to better estimate the rate of drive efficiency decline, we designed and assessed the efficiency of single gRNA drives with imperfect homologous arms, refining a detailed gRNA multiplexing model. To mitigate the greater than expected efficiency loss, we further evaluated two new strategies: (1) extended homology arms to span all target sites with mutations in the PAMs and (2) a population-level multiplexing gRNAs system involving two or more drives, each carrying two gRNAs. Specifically, the population-level multiplexing system has four adjacent gRNA target sites, and the drives have small mutations in their homologous arms to prevent cleavage by the other drive. Mutations in both strategies did not impair efficiency, but they were not consistently inherited, and undesired cutting in the homologous arms decreased drive efficiency. We simulated homing suppression drive using a dual 2-gRNA population-level gRNA multiplexing strategy based on our experimental evaluation. Despite being somewhat more vulnerable to functional resistance than a standard 4-gRNA drive, the higher individual drive efficiency of the population-level multiplexing system increased successful population elimination outcomes. Thus, population-level multiplexing can be a useful for improving suppression drive power.

## Introduction

Gene drive technology is a powerful tool with the potential to address critical challenges across various fields, including public health, agriculture, and conservation, particularly with the application of CRISPR as a genome editing tool^1–4^. CRISPR-guided homing drive can be designed to suppress populations by targeting genes essential genes, or alternatively spread desired traits through genetic modifications within the population^4–7^. This is achieved by spreading the drive allele, which carries the CRISPR system and the desired genetic effect^3,6,8,9^. The drive allele converts wild-type alleles into drive alleles in the germline of heterozygous individuals through homology-directed repair (HDR) following CRISPR-mediated DNA cleavage, thereby increasing its frequency within the population. CRISPR-based gene drives so far have been successfully implemented in a variety of organisms, including yeast^10^, flies^11–13^, mosquitoes^14–18^, plants^19,20^, and mice^21^.

Although the drive allele has the potential to spread throughout an entire population, significant obstacles arise due to end-joining repair mechanisms, which cannot be entirely avoided in CRISPR-based gene drives and must be carefully considered^3,22,23^. End-joining repair introduces mutations or deletions at the target site, rendering it unrecognizable by the gRNA for cleavage. We refer to alleles with mutated target sites as resistance alleles^3^. Resistance alleles have been observed to arise both in germline cells and in the early embryo from maternally deposited Cas9 and gRNA^24,25^. The formation of resistance alleles slow down or even halts the spread of homing gene drive, posing a significant obstacle to its success^18,22,23,26,27^, especially when the alleles preserve target gene function (“functional” resistance alleles). Substantial progress has been made toward avoiding issues with resistance alleles by targeting highly conserved region of essential genes^12,14,28^, suppressing end-joining pathways^29^, and utilizing germline-specific promotors for the expression of Cas9^24,30^. Yet, none of these approaches appear to be sufficient. Single-gRNA suppression drives targeting conserved female fertility genes, such as *doublesex*^31^ or *nudel*^22^ ultimately fail because of functional resistance.

Multiple gRNAs has been suggested as another highly effective strategy to reduce the generation of functional resistance alleles and hence to enable population suppression^7^. Several experimental findings and models back this view, indicating the advantages of applying multiplexed gRNAs^11,23,24,27,32,34,35^. Simultaneous cleavage at two or more target sites is likely lead to a large deletion and loss of gene function^23,34^. Even if cleavage and repair at each site occurs sequentially, it is highly unlikely that all cleavage sites will be repaired in a way that preserves gene function. Nonfunctional resistance alleles can then be eliminated due to their fitness costs^11,28,36^. Lastly, HDR repair can still occur at wild-type gRNA target sites, even if some target sites have already been mutated, which is an additional opportunity for drive conversion^37^. Thus, a second gRNA typically helps improve drive efficiency^11,24^.

However, further increasing the number of gRNAs eventually reduces conversion efficiency due to imperfect homology with the target^35,37–39^. Unless both of the outermost gRNA target sites are cleaved simultaneously, the nonhomologous DNA between the actual cut site and the drive homology arms results in reduced HDR and thus reduced drive conversion efficiency. Two studies examined how such homology mismatch affects the drive efficiency^38,39^. One of them with shorter mismatch reported a notable efficiency decline only when both arms had nonhomologous DNA, but a single arm with extra DNA had no effect^39^. The other study with a longer stretch of nonhomologous DNA reported a notable efficiency drop with one mismatched arm and did not investigate effects of both arms having nonhomologous DNA^38^. This study predicted that because of this effect combined with Cas9 saturation (leading to lower cut rate at individual gRNA target sites as more gRNAs are added), drive conversion rates would decline with more than 2-4 gRNAs^38^. Drive conversion efficiency is one of the key factors that impact the genetic load of a suppression system^40,41^. Genetic load refers to the reduction in the reproductive capacity of a population and serves as an indicator of the drive’s suppressive power. A sufficiently high genetic load is essential for target population elimination^42–44^. Thus, increasing the number of gRNAs to the level necessary to prevent functional resistance may compromise the drive’s ability to completely eliminate or even substantially reduce the target population.

The population-level multiplexing approach was proposed as an alternate multiple gRNA design, separating gRNAs into several independent constructs targeting adjacent regions of the target gene^45^. Several variations of the population-level multiplexing strategy were compared, with the ‘blocking’ strategy demonstrating the best performance. Originally designed for neutral loci, this strategy could also potentially overcome the loss of efficiency of multiplexed gRNAs in the context of a suppression drive, where high efficiency is most critical.

Here, we first investigated in *Drosophila melanogaster* the reduction of conversion efficiency caused by homology discordance in homing drives incorporating either perfect or imperfect homology arms on the left side, right side, or both sides. The measured efficiency reductions were then applied to refine our previous gRNA multiplexing model, and we found that multiple gRNAs performed less efficiently than originally expected, increasing the need for solutions to preserve drive efficiency.

We then evaluated two solutions. First, we designed extensive long homology arms covering all target sites for both sides. The left homology arm is matched to the rightmost target site, and vice versa. This solution, however, only slightly improved performance. Second, we constructed a dual 2-gRNA drive system to assess the concept of population-level multiplexing. The design operates on a principle similar to the “blocking strategy”^45^, but with an extra gRNA for each construct (the optimal number of gRNAs for high performance according to our model). Our experiments showed that the intentional mutations, which were introduced in the homology arm to avoid cleavage from the counterpart drive, were not inherited consistently, allowing undesired cutting that impaired efficiency in the damaged drives. We then modeled a suppression drive that implements the dual 2-gRNA drives design, based on our experimental assessment. Despite this drawback, the population-level multiplexing strategy still retained advantages over conventional gRNA multiplexing approaches, including a superior level of tolerance to functional resistance compared to a 3-gRNA drive and a higher population elimination success rate than 4-gRNA systems over much of our parameter range.

## Methods

### Plasmid construction

The starting plasmids ATSacG, AHDgN4v2, AHDgN1, THDgN1L, TTTg12U2, TTTg34U2, and TTTg1234U4 were constructed in a previous study^38^. Plasmid digests were conducted with restriction enzymes from NEB. PCR was performed with Q5 Hot Start DNA Polymerase (NEB), and DNA oligos were obtained from Xianghongbio DNA Technologies. Gibson assembly of plasmids utilized Assembly Master Mix (NEB), and plasmids were transformed into DH5a competent cells (Vazyme). Plasmids used for injections were purified using the ZymoPure Midiprep kit (Zymo Research). gRNA target sites were found based on the previous research^38^. Detailed drive constructs are shown in Fig. S1. Plasmid sequences of the final drive insertion plasmids and target gene genomic regions with annotation are provided on GitHub (https://github.com/chenwz22/gRNAmultiplexing/).

### Generation of transgenic lines

Lines were transformed by Fungene Transgenic Flies Company. The donor plasmids AHDgN4la, HDgNU4, HDgN12U2, HDgN12U2p, HDgN34U2, HDgN34U2p, THDgN1LRp, THDgN1LR, and THDgN1R were injected separately (350 ng/ul) into the ATSacG line together with TTChsp70c9 (400ng/ul), which provided Cas9 for transformation. AHDgN1 and THDgN1L (previously THDgN 1) lines were from a previous study^38^. Flies were housed in modified Cornell standard cornmeal medium (using 10 g agar instead of 8 g per liter, addition of 5 g soy flour, and without the phosphoric acid) in a 25°C incubator on a 14/10-h day/night cycle at 60% humidity.

### Drive performance assessment

Flies were anesthetized with CO2 and screened for fluorescence using the NIGHTSEA adapter SFA-GR for DsRed and SFA-RB-GO for EGFP. Fluorescent proteins were driven by the 3xP3 promoter for expression and easy visualization in the white eyes of *w*^1118^ flies. Line ATSacG had an EGFP gene located on the chromosome 2L, and all the drive lines in this study were designed to target this EGFP gene. DsRed was used as a marker to indicate the presence of the drive allele. Drive alleles in all lines also had a Cas9 gene regulated by *nanos* elements and a gRNA cassette. Individuals used for assessing drive conversion were produced by crossing ATSacG females (homozygous for EGFP) with homozygous drive males. Drive male progeny were crossed with *w*^1118^ females to evaluate the germline resistance allele formation rate and drive conversion rate (Fig. S2a). Drive female progeny were crossed with EGFP males (line ATSacG) to evaluate the drive conversion rate (Fig. S2b).

### Data analysis

To account for batch effects (each individual cross is considered as a separate batch with different parameters, which could bias rate and error estimates), we analyzed our data as in previous studies^11,38,46^. In brief, fitting a generalized linear mixed-effects model with a binomial distribution (maximum likelihood, Adaptive Gauss-Hermite Quadrature, nAGQ = 25) enables variance between batches, which then results in marginally different parameter estimates but higher standard error estimates. This analysis was performed with R (3.6.1) and supported by packages lme4 (1.1-21) and emmeans (1.4.2).

### gRNA multiplexing model

All stochastic simulations were performed in SLiM (version 4.2.2)^47^, an individual-based population simulation framework. all models share common parameters and ecological components. We used panmictic populations averaging 100,000 individuals with discrete generations. Our multiplexed gRNA model is an updated version of the model from a previous study^38^, considering a single panmictic population of sexually reproducing diploid individuals. The population is defined by the number of male and female adults of each genotype. In each generation, each female randomly selects a male as a mate and then produces progeny for the next generation. Female fecundity is determined by the fitness of the female’s genotype and a density-dependent factor, which is defined as 10/ (1+9 *N*/*K*), where *N* is the current population size and *K* is the carrying capacity. The number of offspring is drawn from a binomial distribution (probability = fecundity/25, max = 50), allowing natural fluctuations around *K*. Each offspring is assigned a random sex, and its genotype is determined by randomly selecting one allele from each parent, with adjustments for drive activity.

The drive allele carries multiple gRNAs, and the wild-type allele contains multiple gRNA target sites. The inheritance of genotypes follows a discrete dynamic process with multiple stages. In this model, we assume that first, early germline resistance alleles form, followed by a homology-directed repair phase (where drive conversion will usually take place) and then a late germline resistance allele formation phase. If the mother of the offspring has at least one drive allele, maternal Cas9 and gRNA can induce additional resistance allele formation in wild-type sites, regardless of which parent they are from. Each resistance allele formation stage is divided into three substages. Wild-type gRNA target sites are first cleaved according to specified cut rates of the phase (adjusted to produce the specified cut rate across all subphases), followed by repair. If multiple sites are cleaved in the same subphase, the DNA segment between the two sites will be deleted when end-joining repair occurs, forming a gap (which always disrupts target gene function). In the early and late germline resistance allele formation phases as well as in the embryo resistance stage, any cleavage is repaired exclusively through end-joining. Only during the HDR phase can drive conversion occur. If HDR fails, the process continues via end-joining repair, generating resistance alleles. Resistance alleles can either preserve or disrupt the function of the target gene. In our model, we assume that 1% of resistance sequences retain target gene function, which is likely a pessimistic assumption for carefully chosen target sites^11,14,24,25^.

The homologous arms of drive are assumed to match to the outermost cut sites. If cleavage occurs at intermediate sites, the intervening DNA will be considered as imperfect homology. The length of the mismatch determines the magnitude of a penalty on HDR efficiency, which multiplies the chance of successful HDR (set to 1 by default). Furthermore, we introduced a ‘Double Mismatch Penalty’ when imperfect homology exists at both the left and right cut sites. This separate penalty also multiplies the chance of successful HDR. When determining the cleavage rate at each gRNA target site in each phase, the model considers Cas9 saturation, which have been shown to be a critical factor influencing the optimal number of gRNAs^38^.

We examined the relationship between gRNA number and drive conversion efficiency, incorporating the double mismatch penalty and updating the repair fidelity coefficient based on experimental data. In our simulations, a wild-type female was crossed with a drive heterozygous male, producing 100,000 offspring. The genotype of each offspring was used to calculate the drive conversion rate. We then examined the effect of gRNA number on population suppression, in which gRNAs are designed to target a haplosufficient female fertility gene. Disrupted null alleles (nonfunctional resistance or drive alleles) result in recessive female sterility. Females with two disrupted alleles are infertile, while males remain unaffected. Drive heterozygotes were introduced at a 1% frequency into a wild-type population of 100,000. We performed 20 replicates for each data point and determined the frequency of population elimination. Simulations were conducted with both the original and the updated models (with harsher mismatch penalty and inclusion of the double mismatch penalty). Three settings were tested: High performance setting: germline cleavage rate = 2%, HDR-phase cleavage rate = 98%, embryo cleavage rate = 5%. Medium efficiency setting: germline cleavage rate = 5%, HDR-phase cleavage rate = 92%, embryo cleavage rate = 10%. Low efficiency setting (though still representing a strong drive): germline cleavage rate = 8%, HDR-phase cleavage rate = 90%, embryo cleavage rate = 15%.

### Population-level gRNA multiplexing model

The population-level multiplexing gRNA model follows the same fundamental process and logic as the standard drive model. Multiple types of initial drive alleles exist, each containing multiple gRNAs, with all gRNA target sites arranged consecutively within the target gene. To simplify the situation for a proof-of-concept, we focused on simulating a dual 2-gRNA scenario, where two types of drive alleles exist, each carrying two gRNAs. Drive allele D_12_ carries gRNAs targeting sites 1 and 2, while drive allele D_34_ carries gRNAs targeting the sites 3 and 4. The initial released drive alleles contain intentional small mutations to make them immune to cleavage at the target sites within the homologous arms. (Fig. 3, Fig. S1). For instance, D_12_ is immune to cleavage by D_34_ and vice versa. However, drive conversion does not guarantee that the newly converted drive alleles can also inherit the intentional mutations in homologous arms. Based on our experimental results, the inheritance rate of intentional mutations was set at 50% (Fig. 3).

When the drive alleles lose their “immunity”, they will become susceptible to gRNA cleavage from the other drive. For instance, if the D_12_ allele loses immunity, the drive is considered as an insertion in between site 1 and 2, while sites 3 and 4, located outside the insertion, are potentially accessible for cleavage. Based on our experimental results if cleavage occurs at these sites, the repair outcomes were assumed to have a 50% probability of a small mutation and a 50% probability of a large deletion between the drive and the outermost site in the homologous arms (Fig. 3). We then implemented a repair fidelity penalty for this specific drive allele by counting the mismatch steps. Thus, the HDR efficiency of the imperfect drive allele will decrease. We similarly simulated a situation with four separate population-level multiplexing drives, each cutting at just one of the four target sites.

Because both drive alleles and wild-type alleles can be cleaved, mismatch steps are calculated by comparing the leftmost and rightmost ends of the two alleles. If a corresponding site has a small mutation, it is counted as 0.5 steps. If a corresponding site is missing due to a gap or deletion, steps are counted based on the number of deleted sites (for example, it is two steps for the 2-gRNA drive that experienced a large deletion). Additionally, it is important to note that if the drive allele and target allele share the same deletion (due to both of them having previously experienced cleavage), it is not counted toward the mismatch value.

Drives are designed to suppress a population by targeting a haplosufficient female fertility gene. We evaluated the drive performance and suppression effects of both population-level gRNA multiplexing (dual 2-gRNAs drives and quad 1-gRNA drives) and standard gRNAs multiplexing (1-gRNA, 2-gRNA, 3-gRNA, and 4-gRNA drives) by varying the functional resistance occurrence rate and the homing phase cut rate. For 1-gRNA, 2-gRNA, 3-gRNA, and 4-gRNA drives, drive heterozygotes were introduced at a 1% frequency into a population of 100,000 individuals. For dual 2-gRNA drives, each of the two drives was released as heterozygotes at an initial frequency of 0.5%, for the same total release size. Quad 1-gRNA drives were similarly each released at a frequency of 0.25%.

## Results

### Assessing repair infidelity induced by imperfect homologous arms

The homology of DNA on either side of a cut site is essential for the efficiency of homology-directed repair. Therefore, one challenge of a multiplexed gRNAs strategy for homing drive is the reduced drive conversion efficiency caused by repair infidelity, as the homologous arms may not perfectly match to each gRNA target site. One study designed a single-gRNA drive to simulate the scenario in a 4-gRNA drive where only the leftmost site is cut and reported a substantial decrease in efficiency^38^. Another study designed three single-gRNA drives to separately simulate cases where the left, right, or both homologous arms with truncated by 20 bp (representing only one mismatching gRNA interval), demonstrating significantly reduced efficiency only when both arms were impaired^39^.

Here, to better assess the impact of unmatched homologous arms on drive conversion efficiency, we designed four one-gRNA drives targeting EGFP, each with homologous arms that matched both, left, right, or neither of the gRNA target sites (Fig. 1a, Fig. S1a). The length of mismatch of each arm was 114 bp, representing approximately the distance from the first to the fourth gRNA. We observed a decrease of 17-38% in drive conversion efficiency when either the left or right homologous arm was mismatched (Fig. 1a, Data Set 1, *p* < 0.01 except male drive with imperfect left arm, z test). There was no significant difference in efficiency between truncated left and right homology arms in males (*p* = 0.595, z test), while a small difference appeared for female drive (*p* = 0.015, z test). The drive with both imperfect homologous ends exhibited a more substantial decrease (58%) in females, and a near-complete loss of drive conversion efficiency was detected in males (Fig. 1a, Data Set 1). A significant difference was detected between the drive with both arms truncated and all other drives for both male and female drive conversion (*p* < 0.001, z test). Moreover, we also tested the germline resistance formation rate in males for all drives, with nearly all EGFP alleles rendered nonfunctional if they were not converted to drive alleles in all systems (Fig. 1a, Data Set 1). Taken together, our results demonstrate that drive conversion is substantially affected by imperfect homologous ends to a greater extent than detected in our previous study^38^, and an additional efficiency reduction when both ends were incomplete^39^ was confirmed and further quantified.

**Fig. 1.**
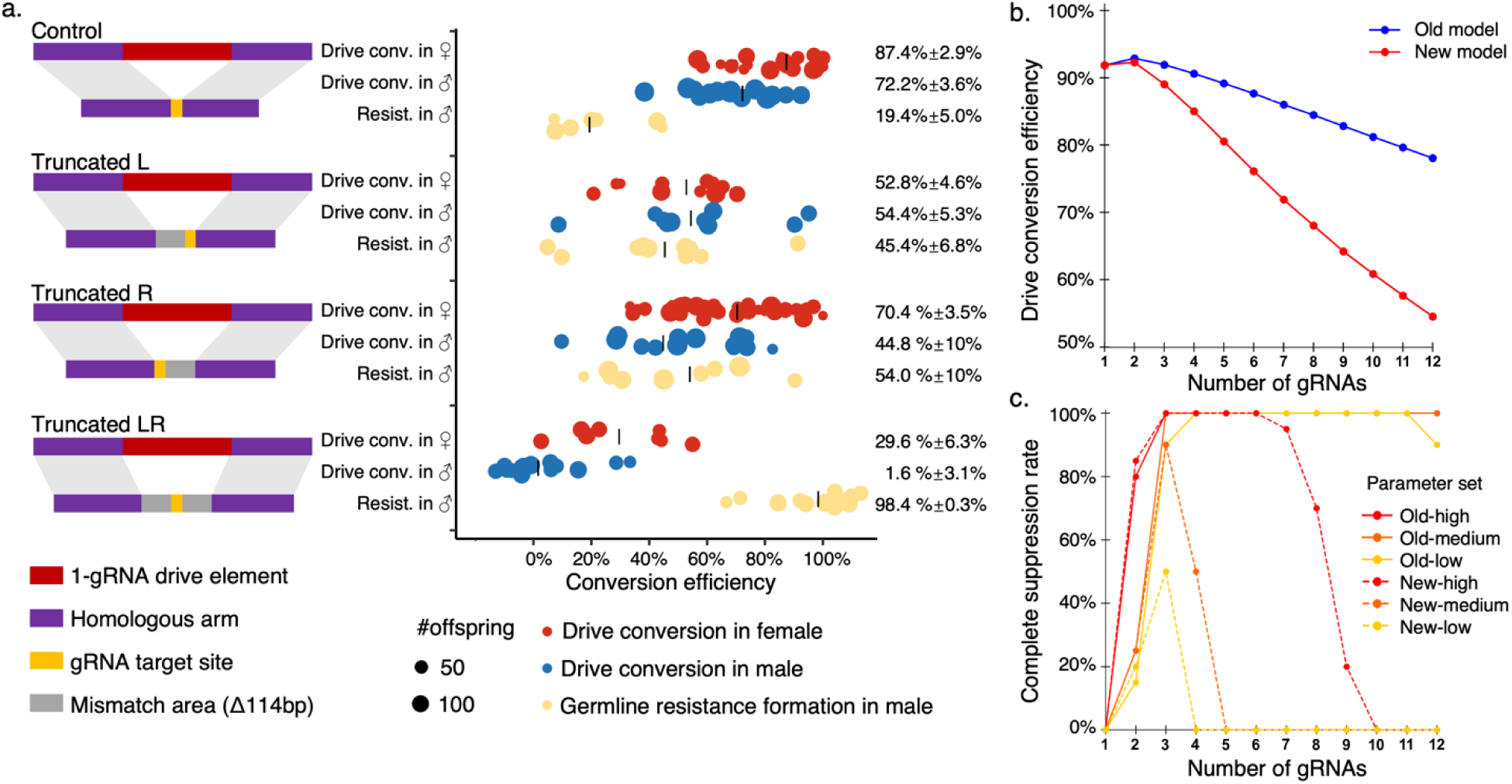
Experimental assessment of drive efficiency and updated gRNA multiplexing model. **(a)** Drive conversion rates and germline resistance formation rates were evaluated in single-gRNA drives with and without truncated homology arms. Each dot represents offspring from a single drive individual. All the drives target the same site within EGFP. The truncation length was 114 bp. Negative drive conversion efficiency indicates that drive inheritance is slightly below 0.5 (no conversion took place). For *p*-values, see Supplemental Data Set 1. **(b)** Using a simulation model, 5,000,000 offspring were generated from crosses between drive/wild-type heterozygotes and wild-type individuals. The graph displays the conversion rate of wild-type alleles to drive alleles. The new model incorporates the double mismatch penalty and an updated (increased) repair infidelity coefficient compared to the old model of a previous study^38^. **(c)** Drive/wild-type heterozygotes carrying a suppression drive were introduced into a population of 100,000 at 1% initial frequency. The displayed results are the fraction of simulations where population elimination occurred. High, medium, and low indicate drive performance (see methods) Each point in both models represents the average of 20 simulations.

### Refinement of the gRNA multiplexing model

The previous study proposed a gRNA multiplexing model to estimate drive efficiency and performance, taking into consideration repair fidelity, Cas9 saturation, and other aspects^38^. The model assumed that the reduction in HDR success rate is correlated to the length of the DNA segment lacking homology (the “mismatch”). The gRNA cut sites are assumed to be equally spaced, so the length lacking homology is indicated by the number of gRNA steps between the outer sites and the actual cleavage site. The penalties from left and right homology mismatches are assumed to be multiplicative.

Using new data, we refined mismatch parameter estimates of the gRNA multiplexing model. Our result demonstrated that drive conversion in the drive with poor homology on either end was about 75% the value of the one-gRNA drive with ideal homology arms. The length of mismatch area is about 114 bp, equivalent to our four-gRNA drive if only the first gRNA cut, thus creating a right arm mismatch of three gRNA “steps” in the drive with a poor right homology arm. Thus, we estimate that the level of repair infidelity parameter to be an 8.3% efficiency reduction per mismatch gRNA step. In cases where both the left and right cut sites exhibit imperfect homology simultaneously, our assessment revealed that the drive conversion efficiency of the drive with poor homology at both ends was approximatelly 14.5% of that observed in the one-gRNA drive with ideal homology arms. We thus introduced a “Double Mismatch Penalty” factor of 0.313, which also multiplies the penalties from left and right homology mismatch to generate the final net HDR penalty.

The previous gRNA multiplexing model showed that the optimal number of gRNAs for maximum drive inheritance is 2-3, perhaps sometimes 4 depending on parameters^38^. Increasing the number of gRNAs would result in moderate reductions in drive efficiency (Fig. 1b). However, our new simulations, which have higher single-mismatch penalties and also have a double mismatch penalty, revealed that drive conversion efficiency is more highly sensitive to the number of gRNAs. The optimal number of gRNAs is now two, and increasing the number beyond this threshold results in a sharper decline in drive conversion efficiency (Fig. 1b).

Suppression homing drives are especially susceptible to failure when drive conversion efficiency is insufficient due to lack of power to eliminate the population, resulting in an equilibrium outcome. We thus assessed how the number of gRNAs could affect the rate at which the drive was successful in eliminating the population under different drive performance regimes (Fig. 1c). The suppression drive targets a haplosufficient but essential female fertility gene. Thus, females carrying two disrupted alleles (drive or nonfunctional resistance) are sterile, while males are unaffected. In the old model, drives with any of our performance settings were usually successful, as long as a sufficient number of gRNAs were present to avoid functional resistance^38^ (Fig. 1c). Only drives with lower performance and a very high number of gRNAs might fail to eliminate the population. However, all drives in the new model became less able to achieve successful suppression, except in a narrow widow with an optimal number of gRNAs for the higher performance systems (Fig. 1c). The population could only be eliminated in all simulations for the high-performance model with four to six gRNAs. While still necessary to avoid failure by functional resistance, when the number of gRNAs exceeds two, the drives rapidly lose suppressive power and quickly become unable to eliminate the target population, even in the absence of functional resistance (though the equilibrium populations are still substantially lower than the original population in these cases). This result indicates that the negative impact of too many gRNAs on homing suppression drive is substantially greater than previously expected.

### Alternate strategy: Long-arm drive alleles with homology arms spanning all gRNA target sites

To prevent imperfect homology from reducing conversion efficiency, we proposed designing extended homologous arms that span all gRNA target sites in the homing drive. In theory, regardless of which target sites are cleaved, the break can reliably align with nearly perfectly matched homologous arms on either side for HDR. We introduced 1 bp mutations for PAM motifs to avoid gRNA target sites in the extended homology ends being cut. The PAM mutations on the left homology arm are differe nt from those on the right arm to reduce repetitive DNA. Ideally, all of each long-arm should be copied during normal HDR, regardless of exactly which target sites are cut.

A significant enhancement in female drive efficiency was observed with the long-arm drive containing extended homology compared to the standard 4-gRNA drive (*p* = 0.043, z test), while male drive efficiency was reduced, though not significantly (Fig. 2, *p* = 0.222, z test). The distribution of drive inheritance among insects with the standard 4-gRNA drive appears bimodal, though the reasons for this are unclear.

**Fig. 2.**
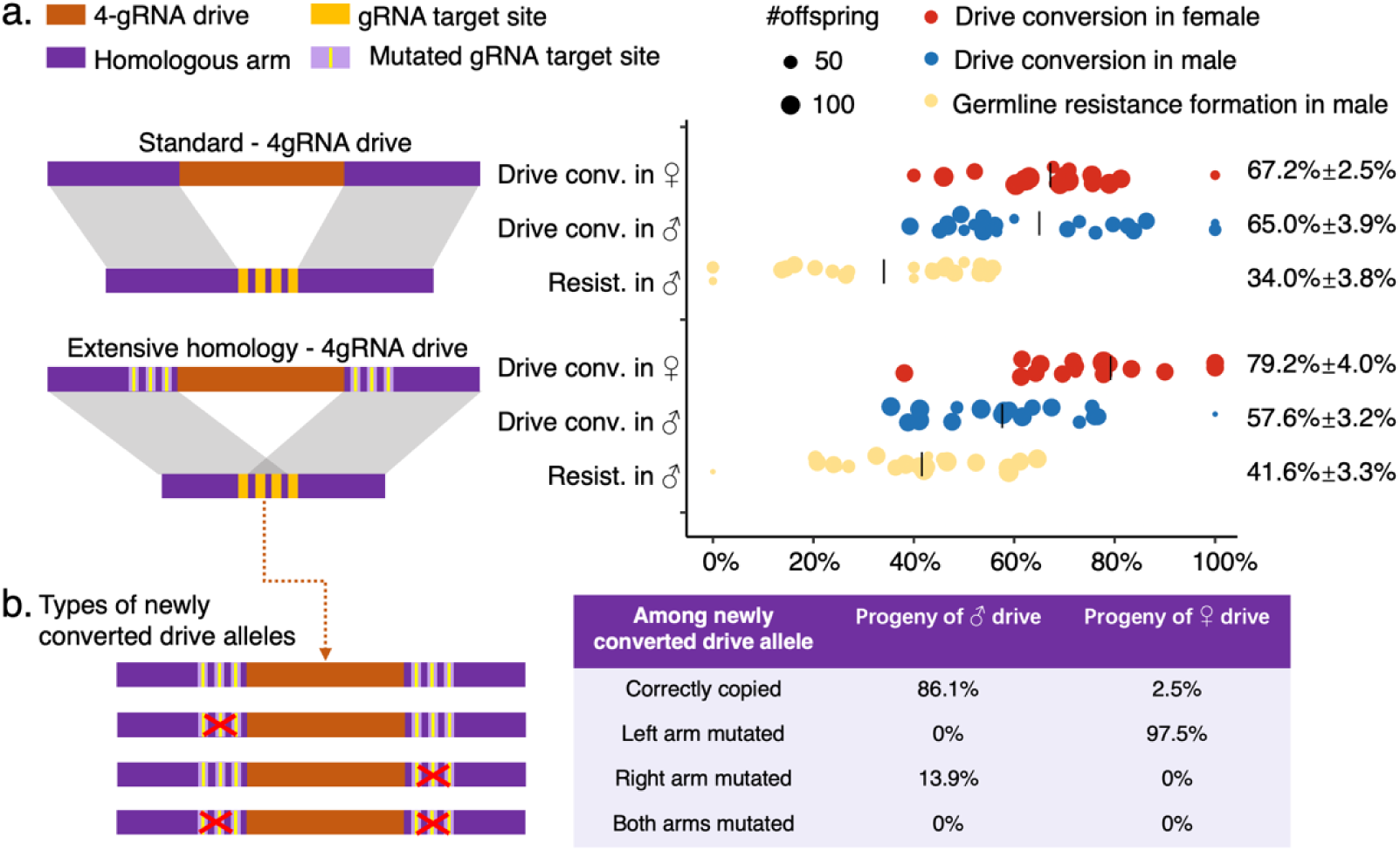
Design and experimental evaluation of the “long-arm” drive. For the standard 4-gRNA drive, its left homology arm matches the cut site of the leftmost gRNA, while its right homology arm matches the rightmost gRNA. For the “long-arm” 4-gRNA drive, its left homology arm is extended to match the cut site of the rightmost gRNA, while its right homology arm matches the leftmost gRNA. PAM motifs in gRNA target sites within extended regions were mutated by 1 bp to prevent gRNA cleavage. **(a)** Drive conversion rates of males and females, as well as the male germline resistance allele formation rate were evaluated. There was no statistically significant difference for male drive conversion between the standard and long-arm drives (*p* = 0.222, z test), while there is significant difference for in females (*p* = 0.043). **(b)** Types of newly converted drive alleles and their frequency are listed, based on the sequence of drive offspring deriving from either male (n=17) or female drive carriers (n=20).

One issue with this drive is that it may lose the long-arm elements or replace them with wild-type DNA. To investigate this, we sequenced drive carrier offspring (Fig. 2) and assumed based on drive performance that a certain fraction were the original drive allele. Of the remainder, likely representing newly converted drive alleles, 13.8% from male drive conversion exhibited mutations in the right homologous arm and none in the left arm. These must have formed by first conversion to wild-type followed by gRNA self-cleavage of the wild-type sites. Among the newly converted alleles from a female drive parent, about 97.5% alleles showed similar mutations in the left homologous arm, but 0% for the right arm. These mutations appeared to have deletions at the gRNA site closest to the drive. However, it remains unclear why there is a large difference in mutation frequency between males and females, as well as why there is an inconsistent preference for mutations in the left versus right homologous arms. This may be due to our limited sample size coupled with the majority of alleles sequenced representing original drive alleles.

We speculate that the long-arms might offer micro-homologous regions for the break site, potentially triggering microhomology-mediated end-joining^48,49^, copying our mutations in the homologous arm to repair cleavage at the target site. This could reduce drive efficiency, though if mutations are carefully chosen, it may not contribute to functional resistance. We also identified potential risks associated with this strategy, specifically, that our designed homologous arms may not be accurately copied into the converted drive allele through HDR. The new drive than immediately suffers from mutations, which will reduce drive conversion efficiency in subsequent generations. It might still be higher compared to a standard drive if the indels are small, but the repetitive DNA on either side of the drive may also reduce efficiency, even if this is likely to be a smaller effect than seen previously^50^ due to limited size of the identity region.

### Alternate strategy: Population-level gRNA multiplexing

The concept of population-level multiplexing gRNAs was first proposed in a modeling study^45^. Rather than releasing one drive construct with multiple gRNAs, this idea was to release multiple drives (each with one gRNA, likely all targeting the same gene) to increase robustness for different performance parameters. Specific alternatives included separate drives, additive drives, overwriting strategy, and blocking strategy. Here, we proposed and tested an improved variant of blocking strategy, based on two gRNAs per drive (Fig. 3a), our predicted optimal number for maximum drive conversion efficiency. For proof-of-concept, we designed two 2-gRNA drives targeting four adjacent gRNA sites. The distance of the outermost cut sites was within 66bp for both drives. We introduced mutations at the PAM motifs of gRNA target sites in the homologous arms for one construct, representing the type of drive that would actually be used in a population-level multiplexing strategy (Fig. 3a). Ideally, this strategy allows each drive to have the highest drive conversion by carrying the optimal number of gRNA, largely escaping the problem of imperfect homology. At the population-level, four sites could still be cut, maintaining functional resistance generation at a sufficiently low level. Alleles resistant to gRNA 1 and 2 can undergo another round of cleavage and homing event when paired with the drives carrying gRNAs 3 and 4 in subsequent generations (Fig. 3a), and vice versa.

**Fig. 3.**
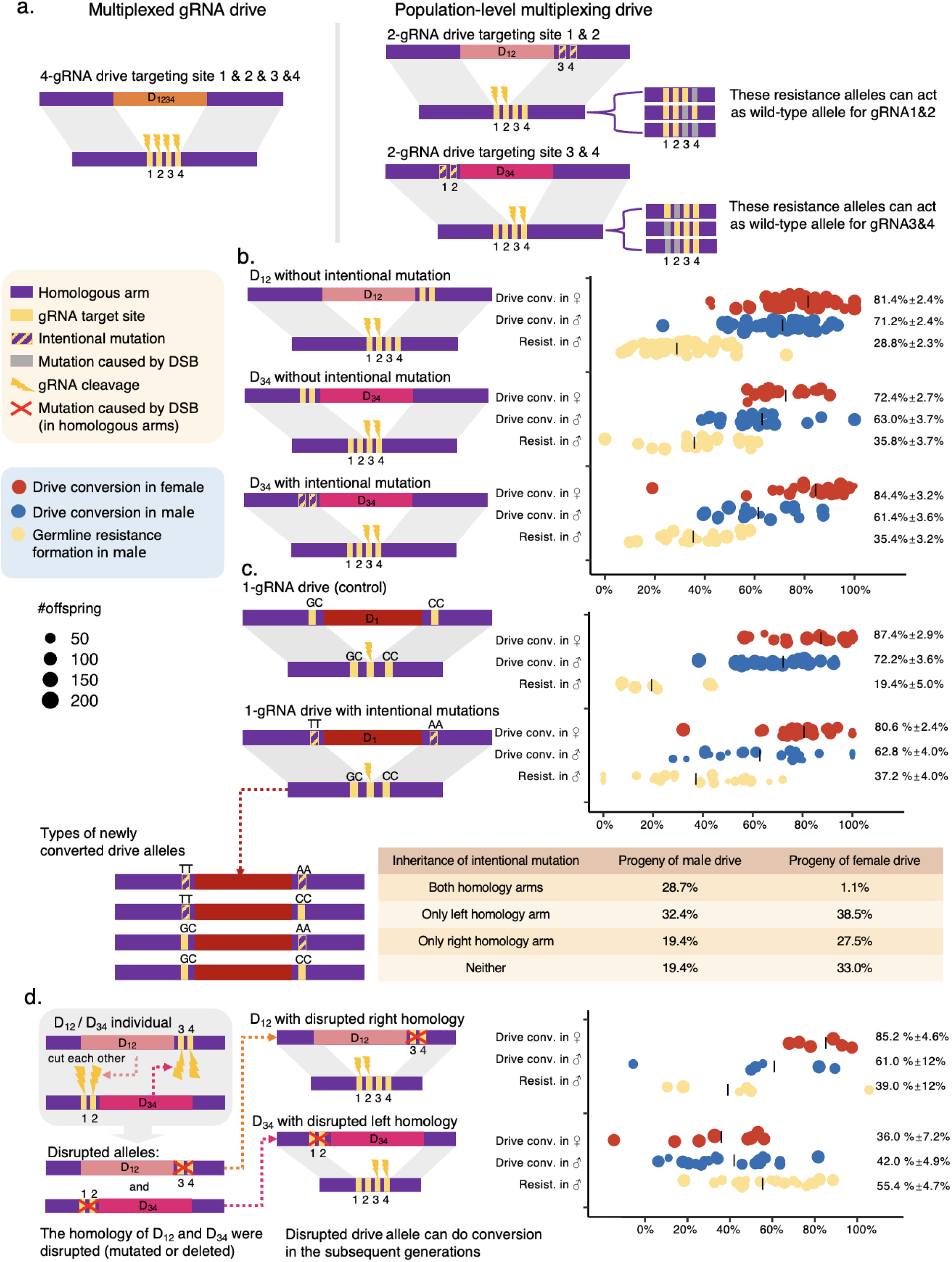
Scheme and performance of the population-level multiplexing drive strategy. **(a)** Concept of population-level multiplexing drive. D_1234_ is a 4-gRNA drive targeting sites 1, 2, 3, and 4 within an EGFP gene on chromosome 2L. D_12_ and D_34_ are 2-gRNA drives targeting sites 1 and 2 and targeting sites 3 and 4, respectively. Because the homology arms are matched to the outmost cut sites for each drive, the PAM motifs in the homology region were mutated by 1 bp in a variant of D_34_ (a population-level multiplexing element) to prevent recognition and cleavage when D_12_ and D_34_ paired with each other. **(b)** Drive conversion efficiency and male germline resistance allele formation were evaluated. No significant differences were observed among the three drives. **(c)** Intentional mutation inheritance in both homology arms was assessed by a 1-gRNA drive D_1_ targeting site 1 and containing intentional mutations within its homologous arms (representing a 1-gRNA population-level multiplexing element in a group of three). The mutated sites were 114 bp away from the drive element. A drive without intentional mutations was used as a control, and no significant difference was found in drive inheritance. However, the intentional mutations within the homology arms were not consistently inherited. Frequencies of newly converted drive alleles are listed, based on the sequence of drive offspring deriving from either male (n=40) or female (n=42) drive carriers. **(d)** Experimental setup where D_12_ is paired with D_34_, each one having sequences that the other could cleave. Drive conversion efficiency of D_12_ and D_34_ were then assessed in the next generation after likely cleavage in one of their homology arms. The conversion rate below zero means that inheritance of the drive was below 50%, indicating no drive conversion.

We first estimated the drive conversion efficiency and resistance allele formation of the drive with gRNAs 1 and 2 (D_12_) and the drive with gRNAs 3 and 4 (D_34_). The drive conversion efficiency was 63-81%, with no significant difference was detected between them (Fig. 3b, *p* = 0.086 for female drive, *p* = 0.061 for male drive, z test). The female drive efficiencies of these 2-gRNA drives were slightlly higher (2-14%) than that of 4-gRNA drive, though only the female drive of D_12_ displayed a significant difference (Fig. 2 and 3b, *p* = 0.003 for female drive of D_12_, *p* > 0.16 for others, z test). Additionally, we evaluated the conversion efficiency of D_34_ with mutated PAMs (Fig. 3c). The results showed that the additional two base pair mutations within the homology did not reduce HDR efficiency. Interestingly, female drive conversion was even slightly increased, though this was borderline and could be due to batch effects (Fig. 3b, *p* = 0.035, z test).

We were concerned that the intentional mutations in the homologous arms might not be precisely copied into a newly formed drive allele. To test this, we constructed a single-gRNA drive with a 2bp mutation in each of its homologous arms (Fig. 3c). The conversion efficiency of this 1-gRNA drive was 63-81%, similar to the control 1-gRNA drive without mutations (Fig. 3c, *p* = 0.15, z test). To determine whether the intentional mutation was inherited in newly converted alleles, we randomly collected drive offspring for genotyping, and factored out the number of original drive alleles likely to be present based on measured drive inheritance. In males, only 29% of newly converted drive alleles inherited the intended mutations from both homologous arms, and in females, this rate was 1.1% (Fig. 3c). The proportion of intentional mutations retained on the right homology arm (19.4% for male, 26% for female) was slightly lower than that on the left arm (32% for male, 39% for female), though this difference was not significant (*p* = 0.762 for male, *p* = 0.804, Fisher’s exact test). Approximately 19% of males and 33% of females inherited no intentional mutations in either homologous arm (Fig. 3c). Additionally, males were more likely than females to inherit the intentional mutation (55% vs. 34%). Compared to the right homology arm, the left homology arm in males also exhibited a slightly higher tendency to successfully incorporate the intentional mutation (50% vs. 39%). However, the inheritance of intentional mutations on the left and right arms was not necessarily correlated. There was no significant difference (*p* > 0.5, Fisher’s exact test), perhaps due to small sample size. This result suggested that the intentionally mutated PAM motifs at gRNA sites within the homologous arms are likely to be lost (about a 44% chance in each conversion) and replaced by the wild-type sequence.

Without the protection of mutated PAMs, the gRNAs expressed by two drive alleles can recognize and cleave each other’s homologous arms when they are paired, introducing mutations or deletions. Drives with shortened or mutated homologous arms may exhibit reduced conversion efficiency. To assess how disrupted homologous arms affect drive performance, we experimentally simulated this scenario. We crossed D_12_ homozygotes with D_34_ homozygotes (both without intentional mutated PAMs) and then crossed their male D_12_/D_34_ heterozygous offspring with *EGFP* homozygotes to separate the two types of drive allele. Due to high cleavage rates, it was assumed that all gRNA target sites would be disrupted, which was supported by our sequencing (see below). We then tested the performance of these drives with disrupted homologous arms (Fig. 3d, Supplement data set 4). No significant difference in drive performance was observed between the original D_12_ and the disrupted D_12_ (*p* = 0.461, z test), whereas a significant reduction was detected for the disrupted D_34_. (*p* < 0.001 for male and *p* < 0.0001 for female drive, z test). The mutation types seen in the homology arms of these disrupted drives varied, including small deletions between the gRNA cut sites and a large deletion of the whole region between the drive itself (not just the inner gRNA target) and the outermost target site (Fig. S3). Notably, we observed that the large deletion of the left homology arm of D_34_ was identical to the arm of D_12_ in 26 out of 29 cases, and the large deletions in the right homology arm of D_12_ matched the right arm of D_34_ in 5 out of 12 cases (Fig. S3), suggesting that the other drive was used as a template for HDR after cleavage. In our designs, the sequences of D_12_ and D_34_ are almost identical except for the extra two gRNA target sites beside the boundary of drive element and the designed homologous arms. The large deletions may have been caused by HDR or perhaps microhomology-mediated end-joining (MMEJ). MMEJ uses small microhomology regions (5–25 base pairs) internal to the break sites for repair of the double-strand break^48,49,51^. These large deletions may have a greater impact on efficiency loss due to the reduced homology in the resulting arms, no longer matching the gRNA target site on one side of the drive.

### Modeling assessment of population-level gRNA multiplexing

gRNAs multiplexing is likely necessary in homing suppression drives to avoid functional resistance, which will could otherwise prevent successful population elimination. However, both standard and population-level multiplexing drives each have potential drawbacks, and it remains unclear if population-level multiplexing represents the superior strategy based on our data. To better understand the possible benefits of this strategy, we simulated homing suppression drives with population-level gRNA multiplexing and compared it to standard gRNA multiplexing drives. We examined variatio ns in the homing phase cut rate (equivalent to the drive conversion rate for the 1-gRNA standard drive in this model) and relative rate of functional resistance alleles (the rate at which a single gRNA target site resistance sequence preserves target gene function when there is no large deletion, which always disrupts function) (Fig. 4). Dual 2-gRNA drives represent our optimal population-level gRNA multiplexing strategy, and we also assessed quadruple 1-gRNA drives (also a population-level multiplexing strategy). The total drive allele release level was held constant (divided into two types of alleles for the dual 2-gRNA population-level multiplexing system, and divided into four types for the quadruple 1-gRNA system).

**Fig. 4.**
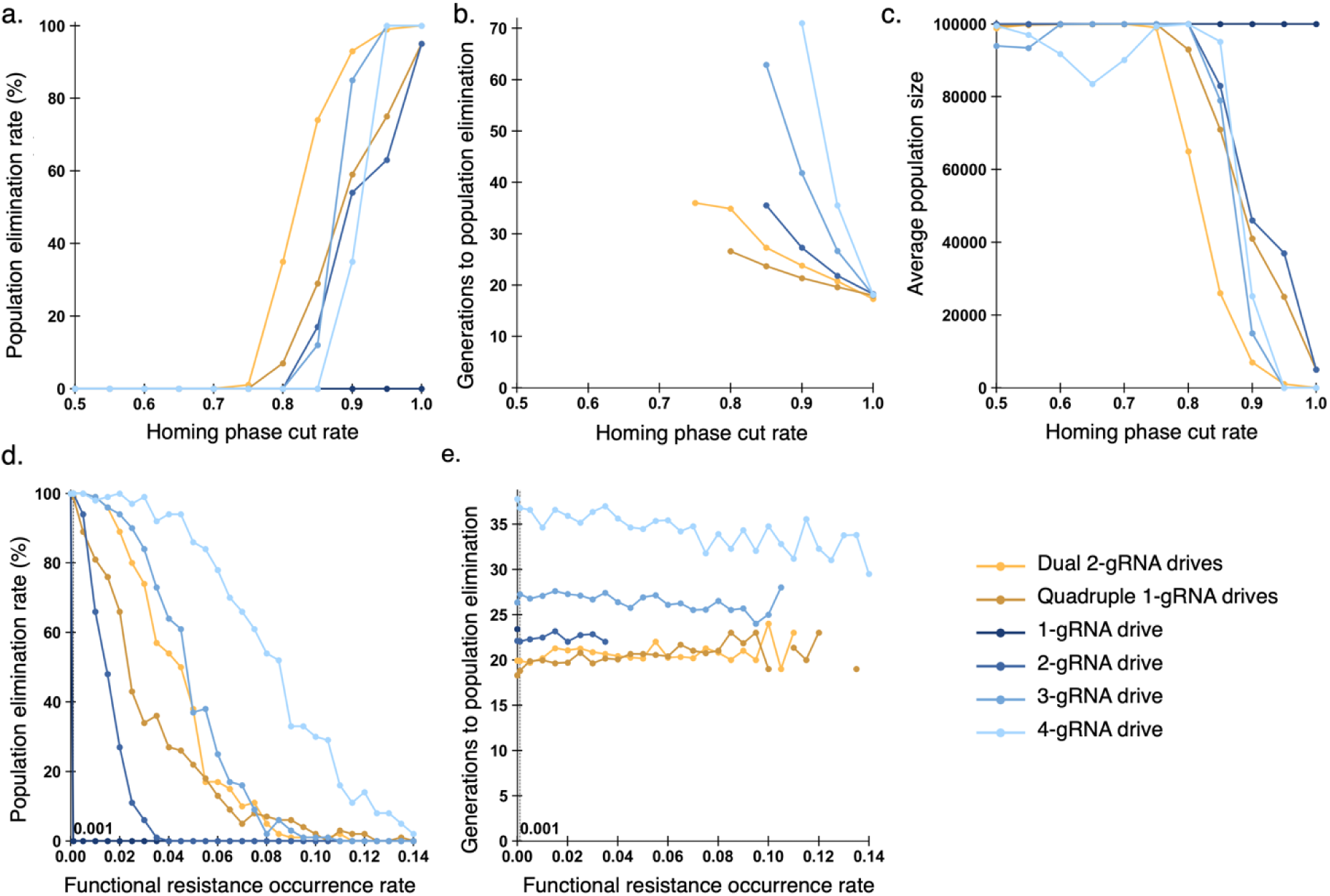
Modeling analysis of standard and population-level multiplexing. Drive heterozygotes carrying a suppression drive were introduced into a population of 100,000 at 1% initial frequency (split evenly between drive types for the population-level multiplexing variants). Simulations were run for 100 generations. Dual 2-gRNA drives and quadruple 1-gRNA drives are designed to target adjacent gRNA sites, representing population-level multiplexing, while the other drives use standard gRNA multiplexing in a single drive. All drives are suppression type targeting a female fertility gene. **(a), (b),** and **(c)** show simulation results under varying homing-phase cleavage rates. 1% of all resistance sequences are functional. Each data point represents the average of 100 replicates. In part c, simulations are extended to obtain a reliable average (population size is reported between generations 300 and 400). **(d)** and **(e)** show outcomes with varying relative functional resistance rate. The homing phase cut rate is 0.95. Each data point represents the average of 100 replicates.

We first conducted simulations with varying homing phase cut rate in the germline stage (Fig. 4 a, b and c). The exact probability of HDR occurrence can be reduced for multiple-gRNA drives or drives with problems in their homology arms (see Methods). The 1-gRNA drive consistently failed to successfully suppress the population due to its sensitivity to functional resistance, despite its high drive conversion rate. With two gRNAs, functional resistance can usually be prevented, and the population can be eliminated if the drive’s efficiency is high enough. However, adding more gRNAs decreases the success rate and increases the required time for population elimination (Fig. 4 a, b and c), especially when the homing phase cut rate is lower. The dual 2-gRNA drive system performs better than standard multiplexed drives under lower homing phase cut rates, achieving a higher success rate and requiring fewer generations for suppression. The quadruple 1-gRNA drive set, on the other hand, is less successful, though when it does succeed, it tends to be slightly faster (Fig. 4 a, b and c). Each drive is immune to cleavage from the other drives at first, but they will gradua **l**y lose this immunity in subsequent generations, which is also when drives meet more often after reaching higher frequency. Thus, the drives cleave each other in males, leading to more homology mismatch that in turns leads to reduced conversion efficiency and decrease of suppressive power.

We further evaluated the tolerance of the gRNA multiplexing systems to functional resistance by varying the relative functional resistance rate over a wide range (Fig. 4 d and e). While we vary these up to a substantially higher level that good target selection would a **l**ow, this may better represent long-term drive performance in very large, natural populations that are beyond our simulation capacity. We found that when drive performance was sufficient to allow population elimination in the absence of functional resistance, the dual 2-gRNA drives exhibited a success rate comparable to that of the standard 3-gRNA drive, indicating a similar level of tolerance to functional resistance. The quadruple 1-gRNA drives performed worse than the standard 3-gRNA drive, but better than the 2-gRNA drive. However, the functional resistance tolerance of both population-level multiplexing drives was substantially weaker than the 4-gRNA drive. This is likely because at low functional resistance rates, population elimination is primarily governed by conversion efficiency, with fewer gRNA target sites keeping efficiency high. However, when the functional resistance rate increases, drive individuals carrying multiple gRNAs have a higher likelihood of generating large deletions due to simultaneous cleavage events, which can substantially reduce the chance of producing functional resistance alleles. Additionally, in population-level multiplexing drives, it is possible for an allele to be functiona l ly resistant to just one drive, giving it a fitness advantage in the population, even if it can still be cleaved by the second drive. This factor likely accounts for the somewhat superior performance per-gRNA for the standard drives over the population-level multiplexing drives. overall, though, both populatio n-level multiplexing drives required the fewest generations to achieve population elimination among all tested configurations. Our model demonstrates that this strategy effectively balances HDR efficiency with tolerance to functional resistance.

## Discussion

Our study shows that the reduced repair fidelity caused by imperfect homology in multiplexed gRNA homing drives is more substantial than previously expected, reducing drive efficiency, which has a large effect on suppression performance. Given that gRNA multiplexing remains essential for reducing functional resistance^38,31,36^, this emphasizes the need for strategies to mitigate this effect. Here, we introduced the long-arm strategy and further explored population-level multiplexing through both modeling and experiments. overall, we found that these methods could potentially allow for improved efficiency, but that they do have complications, preventing fu **l** realization of potential benefits.

Our results show substantially reduced rates of success when both arms have a mismatch (DNA in the cut allele between the cut site and the DNA that is identical to the drive’s homology arms) compared to just one arm, consistent with a previous study^39^. This study reported minimal impact when only one arm was impaired, contrasting with our observations. This discrepancy can be explained by the difference in mismatch length (114 bp in this study compared to only 20 bp in the previous study). Future studies could investigate how the exact length of mismatches in both arms influences repair mechanisms and help further refine our quantitative model.

The experimental assessment of repair efficiency was then used to refine a gRNA multiplexing model for more accurate prediction of homing drive performance. The model was adapted from a previous study, considering Cas9 saturation, timing of cleavage and repair, and other factors^38^. After incorporating more realistic mismatch-related parameters, the updated model shows greater performance loss with increasing number of gRNAs. Once the gRNA number exceeds the optimal value, the successful suppression rate declines significantly more rapidly than in the old model. Even with our best parameter set, the range of gRNA numbers yielding successful population elimination was greatly narrowed. overall, we showed that increasing the number of gRNAs offers only marginal improvements in drive inheritance (the optimal value is now two, when some realistic parameters could previously yield an optimum of 3-4), substantially lower than predicted by previous models^23,32,34,38^. However, multiple gRNAs remain critical for avoiding functional resistance. Our findings provide valuable insights for guiding future experimental design.

First, caution is advised when increasing the number of gRNAs in homing drive designs, according to our modeling result. The gRNA sites should be placed as close together as possible to reduce repair infidelity. Second, the direction and position of gRNAs may matter. D_12_ (two gRNAs with PAMs pointing toward each other) performed slightly better than D_34_ (two gRNAs are in the same direction, with cut sites slightly further apart) (Fig. 3, about 8∼9% difference in conversion efficiency), which can potentially be explained by gRNA activity or the direction of gRNA. Some have argued that mismatch at the PAM-distal end more significantly affects HDR efficiency due to lingering Cas9 binding to the PAM after cleavage, which could explain the discrepancy^39^. Positioning the PAMs of two gRNAs to point toward each other can potentially increase HDR as well by avoiding Cas9 interference post-cleavage.

To mitigate the problem of repair infidelity in standard gRNA multiplexing designs, we evaluated two strategies. The first strategy extended the homologous arms across all target sites, but this did not result in notable improvement in drive conversion efficiency. This may have been because the additional DNA on either side of the drive is repetitive, which has shown to reduce HDR efficiency in a rescue drive^50^. Moreover, based on results from subsequent experiments, we also identified potential issues with this strategy. We found that the intentionally introduced mutations in the PAM motifs in the homology arms could be lost during the conversion process (see below), resulting in the extended homology region being completely identical to the wild-type allele target regions. In this case, the gRNAs expressed by the drive could recognize and cleave its own homologous arms, which may also further interfere with HDR at the same time. Thus, the design of extended homology arms becomes less effective. However, we did not quantify and model these effects, so we cannot rule out that it may still slightly improve the overall system compared to standard multiplexing.

The second strategy was population-level multiplexing^45^. The core idea of this concept is to target multiple sites to prevent functional resistance at the population level, while each individual drive carries only the optimal number of gRNAs for maximization of drive conversion efficiency. Here, we used dual 2-gRNA drives as a typical case to test the population level multiplexing strategy, while also comparing it to the originally proposed population level multiplexing form with quad 1-gRNA drives. Based on our assessment, dual 2-gRNA drives outperformed standard 4-gRNA drives when drive conversion efficiency was the limiting factor, despite targeting the same four sites. Specifically, this strategy should be advantageous in the aspect of drive conversion, and this advantage may be necessary for suppression drives to be successful.

A key aspect of this strategy is the need to introduce specific mutations for PAMs within the homologous arms to avoid two drives cleaving one another. While this would not affect outcomes in females for suppression drives targeting female fertility genes, it could still happen often in males, especially at high drive frequency when drive efficiency is very important for retaining high suppressive power. We confirmed that intentional mutations within the homologous arms do not significantly affect drive efficiency. However, we found that the inheritance rate of these intentional mutations was not high. Understanding the molecular mechanism of HDR sheds light on this. HDR is initiated by the resection of 5′ ends at the DNA break, generating 3′ single-stranded overhangs. These overhangs search for and bind to a homologous template to guide repair synthesis^49,52–55^. There are two aspects that affect the inheritance of intentional mutations. First, the length of resection of the 5′ ends determines the length of donor sequence (drive allele) copied into the recipient allele (wild-type allele)^49,55^. Second, during homolog pairing and invasion, mismatch between invasive 3’ends of the recipient allele and donor sequence can activate the mismatch repair pathway for correction^52^. A previous gene drive study in *Anopheles* showed that mutations in the homology arms of a drive can be transferred along with the drive element^56^. However, in this study, it was somewhat rare for less than 100 nucleotides on either side to be transferred from donor to recipient, while in our study, this was common. This may be due to differences in HDR-related repair pathways in *Anopheles* and *Drosophila*, perhaps related to the higher HDR rates found in *Anopheles* gene drives thus far^57,58^.

In the population-level multiplexing strategy, without the intentional mutations to protect the PAM from recognition, the homologous arm can be cleaved by gRNAs expressed from the other drive. Impaired homologous arms reduce conversion efficiency because they will have mismatch to the cleaved wild-type allele, thereby undermining the advantage of this strategy in enhancing conversion efficiency. Considering the observations and assessment from our experiments, we developed a model to simulate a homing suppression drive based on a population-level multiplexed gRNA strategy. Our model incorporates a detailed calculation of repair fidelity, taking into account the cutting status at each target site as well as potential cleavage sites within the homology arm (e.g. deletions, mutations, intentional mutation), thereby enabling a more accurate simulation of gRNA multiplexing. It may be slightly pessimistic about population-level multiplexing because our parameterization was based on a test where intentional mutations were far from the drive (equivalent to the distance between the first and fourth gRNAs in a 4-gRNA drive, whereas most internal mutations would be closer to the drive and therefore may be preserved at a somewhat higher rate). Even with these imperfections, our result showed advantages in functional resistance tolerance and relatively high efficiency for population suppression. Though the standard gRNA multiplexing strategy is still best at reducing functional resistance, the population-level multiplexing strategy can keep resistance low while boosting drive performance. Additional drives could ensure that functional resistance to all drives is avoided.

Population-level multiplexing still increases complexity, so it is only recommended when suppression drive performance with multiple gRNAs is not already high and when the low-density growth rate or perhaps the chasing propensity of the population is high, requiring additional suppressive power^59–61^. This may in fact encompass a wide array of potential drives because only *Anopheles* drives thus far have demonstrated exceptionally high drive conversion rates. However, this benefit from populatio n-level multiplexing and long-arm strategies may be less critical for modification type drives, which are not substantially affected by modest reductions in efficiency. Furthermore, by splitting the gRNAs into different drives, there is no need to have large numbers of gRNA promoters available, or to risk instability from using the same gRNA promoter for multiple gRNAs in the drive (a particula r ly important consideration because tRNA and ribozyme strategies to express gRNAs from the same promoter may substantially reduce gRNA expression^16^).

Our model holds potential for broader and more in-depth exploration of population-level multiplexing and similar strategies in future studies, yet it should still only be considered as an approximation. More detailed mechanistic studies of drive cleavage and outcomes could further refine the model and perhaps even suggest ways to mitigate problems with this drive system. Even simply more examples of how efficiency changes in various gRNA configurations with actual or simulated partial resistance will allow refinement of performance parameters in the model. Particularly important is determining if penalties to HDR are fairly conserved across species for a given length of mismatch.

overall, this study evaluated the impact of homology discordance on drive conversion, which was subsequently used to refine a gRNA multiplexing model. Two strategies were proposed to improve conversion efficiency. The long-arm strategy appeared less effective, though may still hold some promise, while the population-level multiplexing gRNA strategy, despite some limitations, demonstrated potential for enhancing conversion efficiency and maintaining tolerance to functional resistance, improving outcomes in simulation models. We thus expect that this approach will be a useful strategy for future suppression drive design, especially when the number of available gRNA promoters for a species is limited.

## Supporting information

Supplemental Data Sets

## Acknowledgements

This study was supported by the Center for Life Sciences, the Ge Li and Ning Zhao Life Science Youth Research Fund, and the National Natural Science Foundation of China (grant 32270672). The cluster-based data collection was assisted by High-Performance Computing Platform of the Center for Life Science at Peking University.

## Supplemental Information

**Fig S1.**
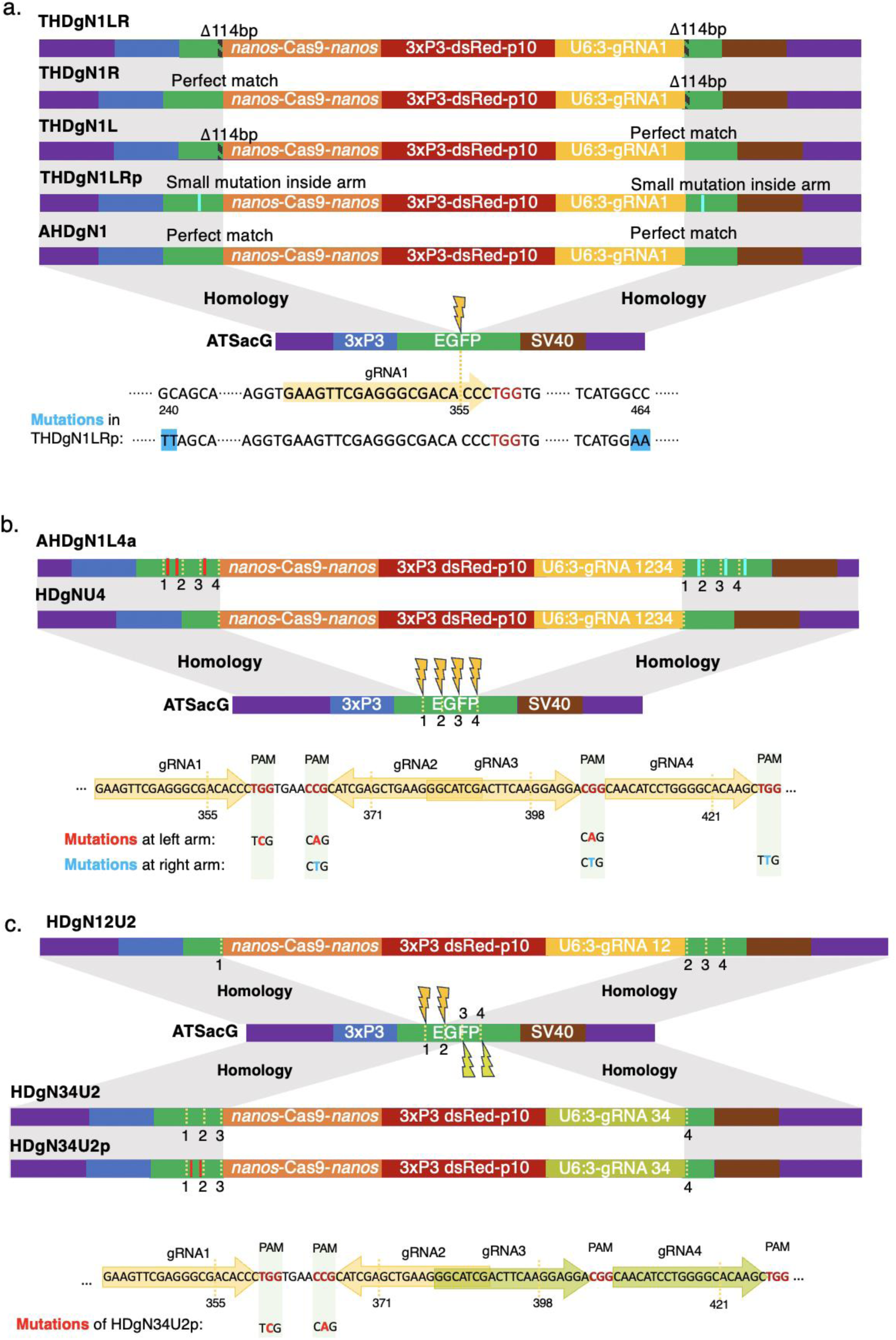
Drive schematic diagrams. The target line, ATSacG, has EGFP driven by the 3xP3 promoter and terminated by the p10 3′ UTR, located on chromosome 2L. The synthetic drive constructs are similar except for the gRNAs and the homologous arms. They contain a Cas9 driven by the germline-specific *nanos* promoter and 3′ UTR, a DsRed marker with a slightly recoded 3xP3 promoter and P10 3′ UTR, and a U6:3 promoter driving one or more tRNA-linked gRNAs that target EGFP. **(a)** AHDgN1, THDgN1L, THDgN1R, and THDgN1LR are designed to assess the repair penalty caused by mismatched homologous arms. AHDgN1 has perfect homology arms for both sides. THDgN1L has a perfect right homology arm and a deletion in the left homology arm, which is 114 bp away from the gRNA cut site. THDgN1R is similar, but with the 114 bp deletion in the right homology arms. THDgN1LR has these deletions in both homology arms. THDgN1LRp is designed for assessing effects in population-level gRNA multiplexing. It is similar to AHDgN1, but with 2bp mutated inside both homology arms. **(b)** Both AHDgN4la and HDgNU4 contain four gRNAs that closely target the EGFP region spanning 355bp-421bp. For HDgNU4, its left homology arm is matched to the cut site of the leftmost gRNA, while its right homology arm is matched to the cut site of the rightmost gRNA. For AHDgN4la, its left homology arm is matched to the cut site of the rightmost gRNA, while its right homology arm is matched to the cut site of the leftmost gRNA. PAM motifs for complete gRNA target sites in the AHDgN4la homology arms were mutated at one bp to prevent recognition and cleavage by the drive’s gRNAs. **(c)** HDgN12U2, HDgN34U2, HDgN34U2p are designed for assessing the population-level multiplexing. HDgN12U2 carries gRNA 1 and 2, while HDgN34U2 and HDgN34U2p carry gRNA3 and 4. Their homologous arms are matched to the cut site of their outermost gRNAs. For HDgN34U2p, the PAM sequences of the gRNA1 and gRNA2 target sites inside the left homology arm are mutated.

**Figure S2.**
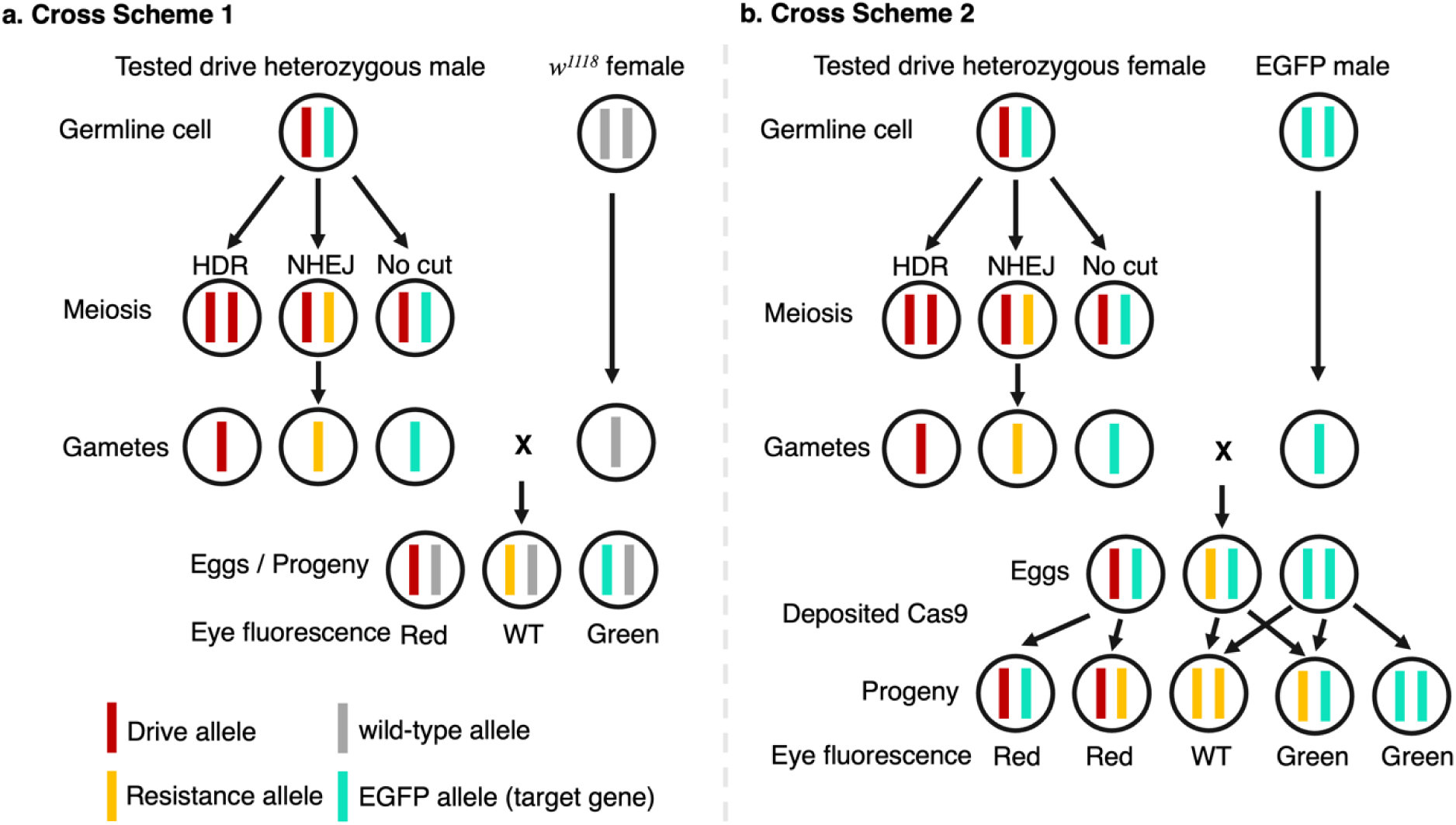
Illustration of cross schemes. **(a)** Drive heterozygous males were crossed with *w^11^*^18^ females. Drive conversion in males can be inferred from the proportion of drive-carrying progeny. Wild-type offspring indicate mutations at the EGFP target site. Therefore, the fraction of wild-type progeny can be used to estimate the germline resistance allele formation rate in males. (**b)** Drive heterozygous females were crossed with EGFP homozygous males. Drive conversion in females can be inferred from the proportion of drive-carrying progeny. Embryo resistance formation and mosaicism can also be assessed.

**Figure S3.**
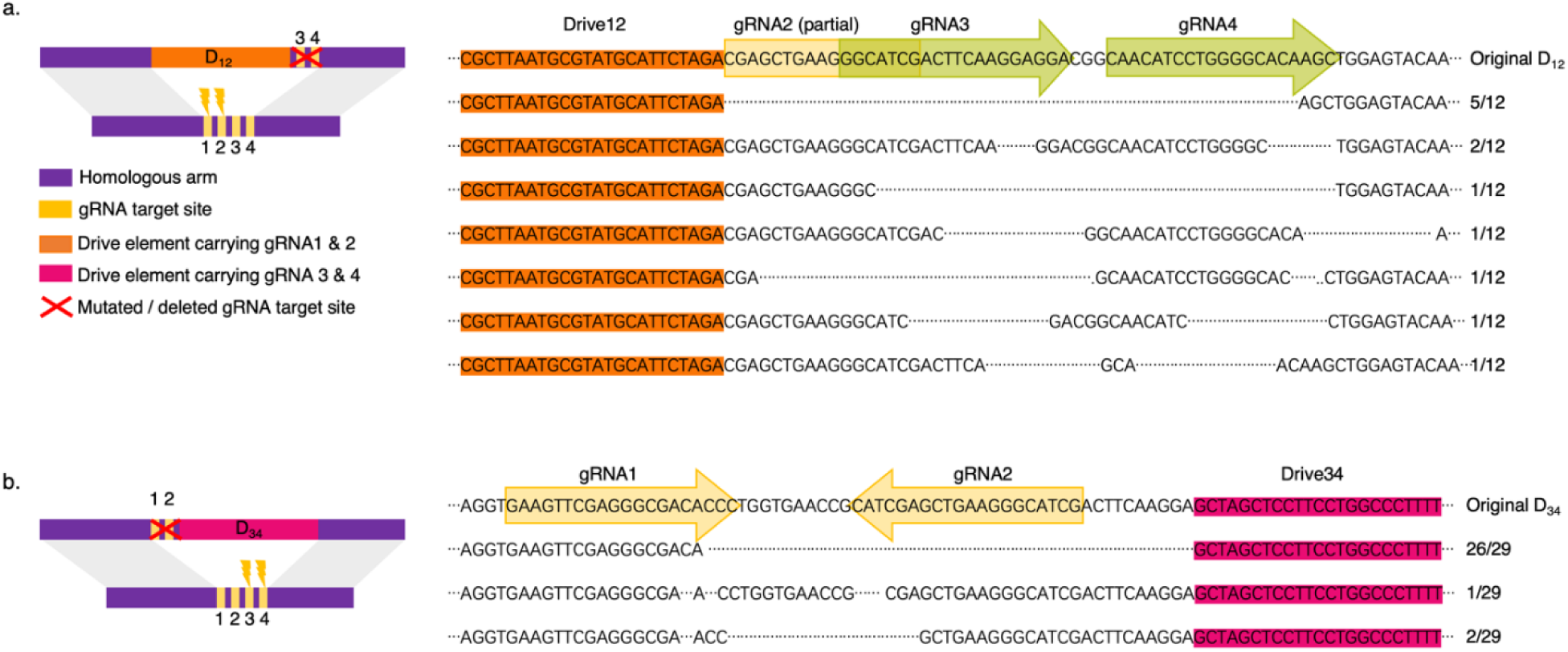
Mutation types at target sites within homologous arms of D_12_ and D_34_ drives. The disrupted drive carriers were the offspring of D_12_/D_34_ drive males. Without the intentional mutations, the drive alleles cleaved each other’s homologous arms. The sequence and the proportion of each mutation are shown. **(a)** Mutations in the right homologous arm of D_12_. **(b)** Mutations in the left homologous arm of D_34_.

## Notes

https://github.com/chenwz22/gRNAmultiplexing/

### Competing Interest Statement

The authors have declared no competing interest.

